# Commentary on Sanborn and Chater: Posterior Modes are Attractor Basins

**DOI:** 10.1101/109355

**Authors:** Phillip M. Alday, Matthias Schlesewsky, Ina Bornkessel-Schlesewsky

## Abstract

Sanborn and Chater [1] propose an interesting theory of cognitive and brain function based on Bayesian sampling instead of asymptotic Bayesian inference. Their proposal unifies many current observations and models and, in spite of focusing primarily on cognitive phenomena, their work provides a springboard for unifying several proposed theories of brain function. It has the potential to serve as a bridge between three influential overarching current theories of cognitive and brain function: Bayesian models, Friston’s [2–4] theory of cortical responses based on the free-energy principle, and attractor-basin dynamics [5,6]. Specifically, their proposal suggests a high-level perspective on Friston’s theory, which in turn proposes a sampling procedure including appropriate handling of autocorrelation as well as a plausible neurobiological implementation. In turn, these two theories together link into attractor-basin dynamics at the level of networks (via Friston) as well at the level of behavior (via the relationship between the modes of prior and posterior distributions, as discussed by Sanborn and Chater). We will argue here that, by linking Sanborn and Chater’s approach to neurobiological models based on the free-energy principle on the one hand and attractor-basin dynamics on the other, the scope of their proposal can be broadened considerably. Moreover, a unified perspective along these lines provides an elegant solution to several of Sanborn and Chater’s Outstanding Questions relating to the neural implementation of sampling.

---

Sanborn and Chater briefly touch upon the connection of their work to Friston’s hierarchical model proposal, but only link it rather generally to the broad computational approach he has proposed for the representation and computation of these models in neural wetware [7]. Friston’s other work, however, also describes and models the relationship between behavior and the sampling procedure necessary for active Bayesian inference [8]. This is compatible with the phenomena that Sanborn and Chater describe as ‘warm-up’ tuning. Although Sanborn and Chater perhaps intentionally formulated their proposal in an implementation agnostic way, Friston’s approach fills in the gaps regarding the neural implementation of sampling in an illuminating way that has been used to model a wide range of phenomena [3,4,8,9]. In particular, Friston’s model provides large-scale suggestions – at the level of groups and networks of neurons – of how sampling is implemented (i.e. hierarchical structure [2,7] and active sampling [2]), and suggests that a simplified or indirect probability distribution is used, i.e. free energy as a proxy for model evidence [8]. In this framework, autocorrelation is minimized via the active sampling procedure, but is also effectively handled by the iterative model updates – autocorrelated samples provide little additional information and thus small error signals. They therefore contribute in decreasing amounts to overall model convergence (see Figure 1). This is related to the performance of particle (i.e. Kalman) filters, which Sanborn and Chater also mention.

**Figure 1:**
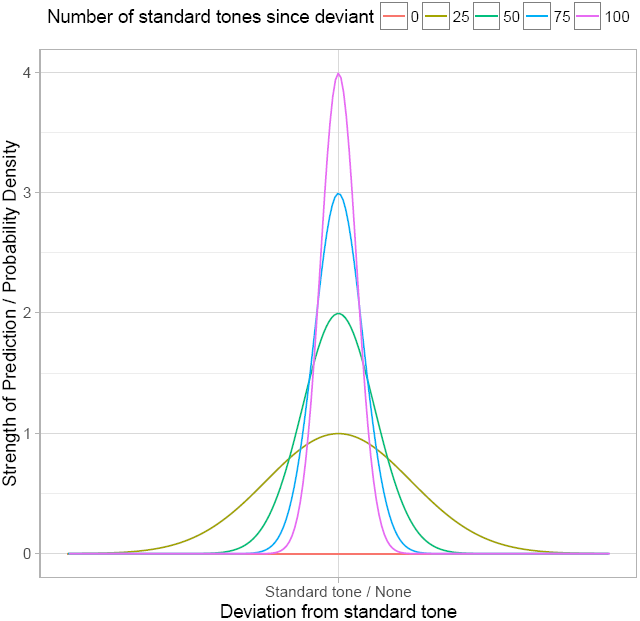
Posterior density of predictions in an auditory oddball paradigm. In Friston’s mismatch negativity (MMN) example [2,9], the MMN reflects the prediction error; in other words, it correlates negatively with the posterior distribution resulting from the previous tone. Autocorrelation of samples is not a large contributor in this iterative framework, because each additional standard tone contributes little new information and the rate of convergence drops off as the previous model state approaches the actual input, seen here in the changing width of the modal peak, despite continued rapid strengthening of an individual point prediction (the height of the modal peak). In Bayesian terms, the posterior does not differ much from the prior when the model evidence does not differ much from the prior. For example, the difference between a 90-10 match-mismatch and a 99-1 ratio is much smaller than that between a 70-30 and a 61-39 ratio, even though the prediction has become stronger (seen here in the height of the modal peak). Attractor basins arise when posterior beliefs from one model iteration are used as prior beliefs for new model (and can be visualized by inverting the density plot). Values here are schematic and calculated from a normal distribution with precision (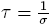) equal to number of tones divided by 10.

While Friston’s model has been proposed as an overarching theory describing the brain as a whole, attractor basins have been proposed as an explanation of emergent classifier behavior in dynamical systems such as neural networks (e.g. [5,6]). Attractor basins are steady states in a dynamical system that are associated with stable, high-firing rates in neural networks, whether computational or biological. In many ways, this approach complements Friston’s principles-first approach with an emergent, empirical observation, yet attempts to connect these two theories thus far have been restricted to observations in passing such as attractors as local optima (and hence stable states) in the free energy landscape, without regard to their concrete implementation or emergence during the sampling procedure in Friston’s model. Sanborn and Chater’s approach provides exactly this connection because the posterior modes in their sampling procedure are essentially attractor basins – areas of high probability density where a posterior belief tends to be drawn and to which estimates (beliefs) tend to converge (see Figure 1). This observation goes beyond Sanborn and Chater’s connection to the mechanisms of neural networks such as the Boltzmann machine and deep belief networks and underscores the deep relationship between these two theories. The Bayesian combination of prior beliefs (including modes / pre-existing attractor basins) and the likelihood (model based on current evidence) leading to a new set of modes when the evidence is strong enough but subject to bias from finite sampling also provides a convenient explanation for the emergence of new attractor basins, i.e. perceptual categories and decisions, over time. As attractor networks provide a neurobiologically plausible way of modelling neural processes related to decision making and classification across a range of scales from perception [10,11] to more complex cognitive domains such as language processing [12], this isomorphism – both in sampling behavior (as noted by Sanborn and Chater) and in large scale behavior via sampling modes and attractor basins – is particularly interesting.

Sanborn and Chater’s proposal thus provides deep connections to leading theories of neural organization and their emergent dynamical and behavioral properties. Given its direction application to cognitive phenomena, their proposal thus provides a potential missing link in a “trinity” of models of the brain and behavior from the lowest levels of organisation (small-scale networks) all the way to its highest levels (cognition and behavior).

